# Antibiotic tolerance is associated with a broad and complex transcriptional response in *E. coli*

**DOI:** 10.1101/2020.08.27.270272

**Authors:** Heather S. Deter, Tahmina Hossain, Nicholas C. Butzin

## Abstract

Antibiotic treatment kills a large portion of a population, while a small, tolerant subpopulation survives. Tolerant bacteria disrupt antibiotic efficacy and increase the likelihood that a population gains antibiotic resistance, a growing health concern. We examined how *E. coli* transcriptional networks changed in response to lethal ampicillin concentrations. We are the first to apply transcriptional regulatory network (TRN) analysis to antibiotic tolerance by leveraging existing knowledge and our transcriptional data. TRN analysis shows that gene expression changes specific to ampicillin treatment are likely caused by specific sigma and transcription factors typically regulated by proteolysis. These results demonstrate that to survive lethal concentration of ampicillin specific regulatory proteins change activity and cause a coordinated transcriptional response that leverages multiple gene systems.

## Introduction

When disaster strikes, a coordinated response is required to mitigate the effects and survive in the face of adversity. Cells have developed an array of regulatory networks to coordinate cellular activity and respond to their environment. These regulatory events occur simultaneously and at multiple levels, including transcriptional, translational, and post-translational regulation. The identification of individual regulatory interactions in combination with bioinformatics approaches has resulted in multiple databases detailing regulation throughout the cell, the most detailed databases are currently for *E. coli* (e.g. RegulonDB^2^, EcoCyc^3^). The depth and availability of information regarding regulatory dynamics provide a new framework for analyzing sequencing data in the context of the whole cell. As such, these types of databases have been applied to analyze network topology^4^ as well as experimental datasets^5–9^. This data has been used to examine transcriptional regulatory networks (TRNs) at the transcriptomic level, which consider gene expression as a function of regulatory interactions by DNA-binding proteins (transcription factors and sigma factors)^10^. We focused on RNA-sequencing data in the context of cell growth and antibiotic stress. Antibiotics disrupt microbial systems and cause targeted cell death; bacterial survival of antibiotics is therefore dependent on cellular networks adjusting to circumvent the physiological impact of the antibiotic.

Antibiotic tolerance, and more specifically antibiotic persisters, are a perfect example of how alterations in metabolic activity (i.e. gene expression and protein activity) lead to changes in cell phenotypes, increasing the chance of survival. Persisters are a subpopulation of cells that tolerate antibiotics for long periods of time despite being genetically identical to susceptible cells. After antibiotic treatment, the surviving population may mutate or acquire resistance genes from their environment^11–13^. Persistence and antibiotic tolerance have been studied extensively, but the heterogeneity of the tolerant population combined with the variance between antibiotic survival of diverse species and growth conditions has led to a number of different, often conflicting, models regarding the causative factors underlying antibiotic tolerance. Some conflicting reports suggest the presence of subpopulations that are metabolically repressed, particularly with respect to translation, while others have found that active metabolism plays a vital role in antibiotic tolerance^14–18^. A few studies have leveraged ^19–21^RNA-sequencing (RNA-seq)^18,22–23^ and proteomic data^24–26^, and they have identified several individual genes and systems related to antibiotic tolerance.

Other work has used gene knockouts and/or overexpression to study specific elements of antibiotic survival^27–29^. However, knockouts and gene overexpression can lead to disrupted cellular networks, alter growth rates, and impact mechanisms that alter awakening from dormancy (growth rates and dormancy have been well-studied^30^). Nonetheless, these works and others have attributed several different mechanisms to antibiotic tolerance. Many genes identified by these studies are sigma factors or transcription factors, regulatory proteins that affect gene expression^26,29,31–33^. The significance of these genes to antibiotic survival suggests that a broader regulatory network underlies the antibiotic stress response. Although understanding the role of individual genes in antibiotic survival is important, what is currently lacking is a more holistic understanding at the network level. While it has been a long-standing goal to identify one or more genes unique to antibiotic tolerance, our TRN analysis shows that the transcriptional response to antibiotic treatment is controlled via a whole-cell, network response consisting of smaller, overlapping networks, some of which are redundant.

We have taken a systems biology, top-down approach to explore antibiotic survival of tolerant populations, and obtained deep, RNA-seq data on both antibiotic-treated and -untreated populations emerging from stationary phase. Our results show that cells have an active and coordinated response to lethal antibiotic stress that alters the activity of specific sigma factors and transcription factors (TFs). Existing literature allows us to connect the activity of these regulatory proteins and changes in gene expression to the pathways involved in antibiotic tolerance. Our findings suggest that many of the regulatory proteins activated during antibiotic stress are regulated by protein degradation. Protein aggregation was recently demonstrated in persister cells^26^, and our previous work shows that interfering with degradation at the protease ClpXP increases antibiotic tolerance in *E. coli*^*34*^. To test if the transcriptional changes are due to altered proteolysis, we then compared the transcriptome of populations with and without slowed protein degradation using our previously established system^34^. The analysis of the TRNs in these populations provides evidence that slowed proteolytic processing of native proteins activates a stress response that promotes recovery once the stress is removed.

## Results and Discussion

We hypothesize that bacteria survive antibiotics by a network response where intracellular networks communicate to adjust cellular responses, rather than a single gene or system as the sole agent of antibiotic tolerance. While individual genes or systems can alter a population’s ability to survive antibiotics, we assert that the individual genes/systems regulate the response but that multiple pathways play a role in antibiotic survival. Here, we support this hypothesis using transcriptional data from antibiotic-treated and untreated populations, and then compare our results to the current knowledge in the field. We attest that the conflicting data surrounding genes related to antibiotic survival (persistence and tolerance) in the literature is due, in part, to researchers slightly or significantly altering network dynamics through gene knockouts and overexpression, and that a more holistic system biology approach could provide new insights.

### RNA-sequencing of ampicillin treated and untreated cultures

We will initially discuss gene expression in three different populations: *a stationary phase population*, which was then diluted into fresh media and incubated for three hours without ampicillin (*untreated*, Amp-) or with lethal concentrations of ampicillin (*amp-treated*, Amp+). We used rapid filtration to harvest cell samples followed by flash freezing in liquid nitrogen. This allows us to recover any whole cells present after antibiotic treatment regardless of viability (this includes persisters and cells that are considered “viable but not culturable” (VBNC)). An advantage to this method is that it enables the concentration of cells while minimizing the possible time for RNA degradation. We found that rapid filtration has greater than 2.5 times fewer dead cells during harvesting than pelleting through centrifugation (a commonly used alternative to filtration). The VBNC levels were similar in the untreated and treated populations; we observed that VBNCs levels were 2.7-fold and 2.2-fold higher than culturable cells (counted on agar plates) in stationary phase and ampicillin treated cultures respectively. Recent research has suggested that VBNC cells are physiologically related to persisters; both populations are often found together^35^, increase due to stress^36–37^, appear stochastically in non-stressed environments^38^, and can withstand greater stress than the susceptible population^35,39–42^. However, the presence of VBNCs demonstrates the heterogeneity of the population that survive antibiotic treatment, but VBNCs are difficult to separate from persister cells. Regardless, we are interested in the tolerant population as a whole and the systems we have identified relating to antibiotic tolerance may be relevant to the whole surviving population or specific subpopulations.

This work builds off our previously reported results with antibiotic tolerance, and uses a similar setup with the same strains of *E. coli*^*34*^. The sequencing data was analyzed using DESeq. Here we consider significant changes in gene expression to be greater than two-fold with an adjusted p-value <0.1. We began by examining changes in gene expression between the stationary phase and the untreated populations. Comparing these two populations allows us to identify how the cells adjust to their new environment, as diluting the cells into fresh media is a critical aspect of selection for ampicillin tolerance from stationary phase (ampicillin is most effective on growing cells)^30,43–44^. The majority of the genes with significantly different expression in the untreated population decreased from stationary phase (rather than increased), suggesting that cells are expending fewer resources in the new media. This phenomenon is likely due to the introduction of fresh nutrients and dilution of spent media resulting in a less stressful environment. Data for the untreated population provides information on the changes specific to the new environment; thus, we can focus on changes specific to antibiotic stress by identifying differences in gene expression unique to the treated population. These differences provide key insights into the molecular mechanisms of antibiotic tolerance and the role of transcriptional regulatory networks (TRNs) in antibiotic survival and recovery.

### Clusters of Orthologous Groups (COGs) during ampicillin treatment

Using a top-down approach, we analyzed changes in gene expression based on the general functional classification, as defined by Clusters of Orthologous Groups (COGs)^1,45^. Although the COG classification system is limited to known functions, more qualitative than quantitative, and often used to study evolutionary relationships; it provides a practical method of grouping genes to identify notable trends in the data. When we analyze the COGs for gene expression unique to the amp-treated population, we find that a few categories stand out (Fig. 1). We will focus on groups with known functions; however, there are seven genes categorized as “function unknown” or “general function prediction only” upregulated during ampicillin treatment, which suggests their function is related to survival or recovery (Fig. 1).

**Fig. 1.**
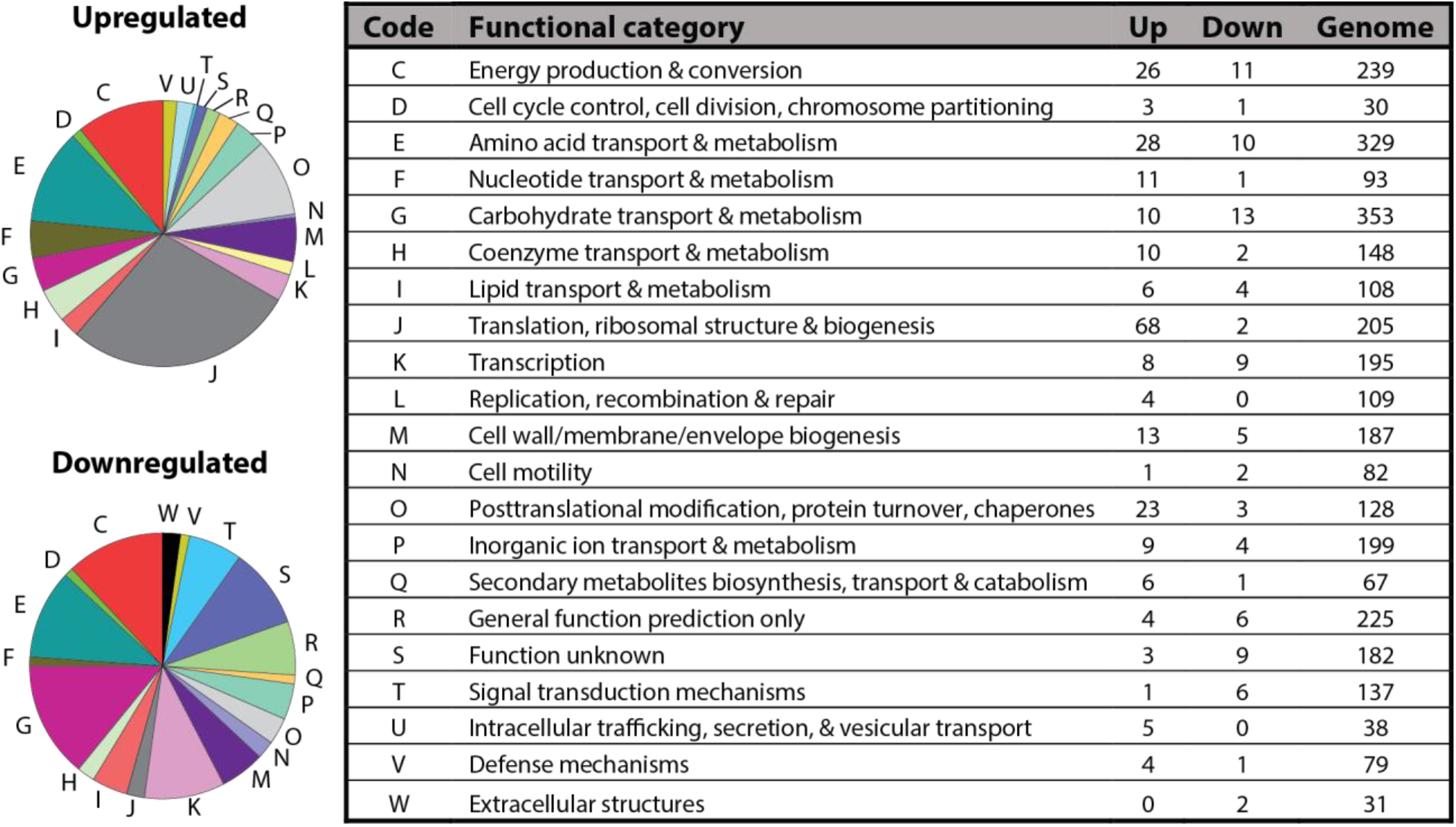
Cluster of Orthologous Genes (COGs). Changes in gene expression specific to ampicillin treatment are enriched in a few COG categories but are rare or nonexistent in others. Left: The number of genes (counts) that are differently expressed (adjusted p<0.1) in the ampicillin treated population organized according to COGs (Top left: upregulated; Bottom left: downregulated). Only genes that were unique to the treated population (compared to stationary phase) were included; genes that show differential regulation in both the treated and untreated population (compared to stationary phase) were removed. Genes in multiple functional categories were accounted for more than once. Right: Functional categories and codes from REF^1^. Up: increased >2-fold gene expression. Down: decreased <2-fold gene expression. Genome: number of genes with that functional category based on the annotated genome.

The most upregulated category is translation, ribosomal structure and biogenesis (68 genes). While the accumulation of proteins involved in translational machinery has previously been shown in ampicillin treated *E. coli*^26^, it is important to note that we removed rRNA during the RNA-seq library preparation process and most of the genes in this category encode ribosomal proteins. Thus, we will not go into depth for this particular group to avoid potential biases in our datasets. The remaining groups with more than 20 upregulated genes are amino acid transport and metabolism (28 genes); energy production and conversion (26 genes); and posttranslational modification, protein turnover, chaperones (23 genes). We explore these groups further in the context of the TRN.

### Analysis of the sigma factor (σ) regulatory network shows that the activity of σ^70^, σ^38^, and σ^32^ increase during ampicillin treatment

Sigma factors are the master regulators of gene expression and essential components of transcriptional regulation. These proteins interact with RNA polymerase to initiate transcription, and their activity changes depending on cellular conditions^46^. Sigma factors are often regulated at the post-translational level by anti-sigma factors or alterations in protein degradation, and different sigma factors are active under specific circumstances. The activity of some sigma factors can be altered by environmental conditions with little change in their mRNA level^47^. However, we can contextualize changes in gene expression controlled by sigma factors by using TRN analysis. To do so, we used all the available information on the direct regulation of all annotated genes from EcoCyc^3^ as defined at the time of analysis (File S1). We compared overall transcriptional activity by mapping changes in expression to specific sigma factors in the context of COGs to get a global picture of gene expression (Fig. 2). We are the first (to our knowledge) to use COGs to support a TRN analysis based on experimental data.

**Fig. 2.**
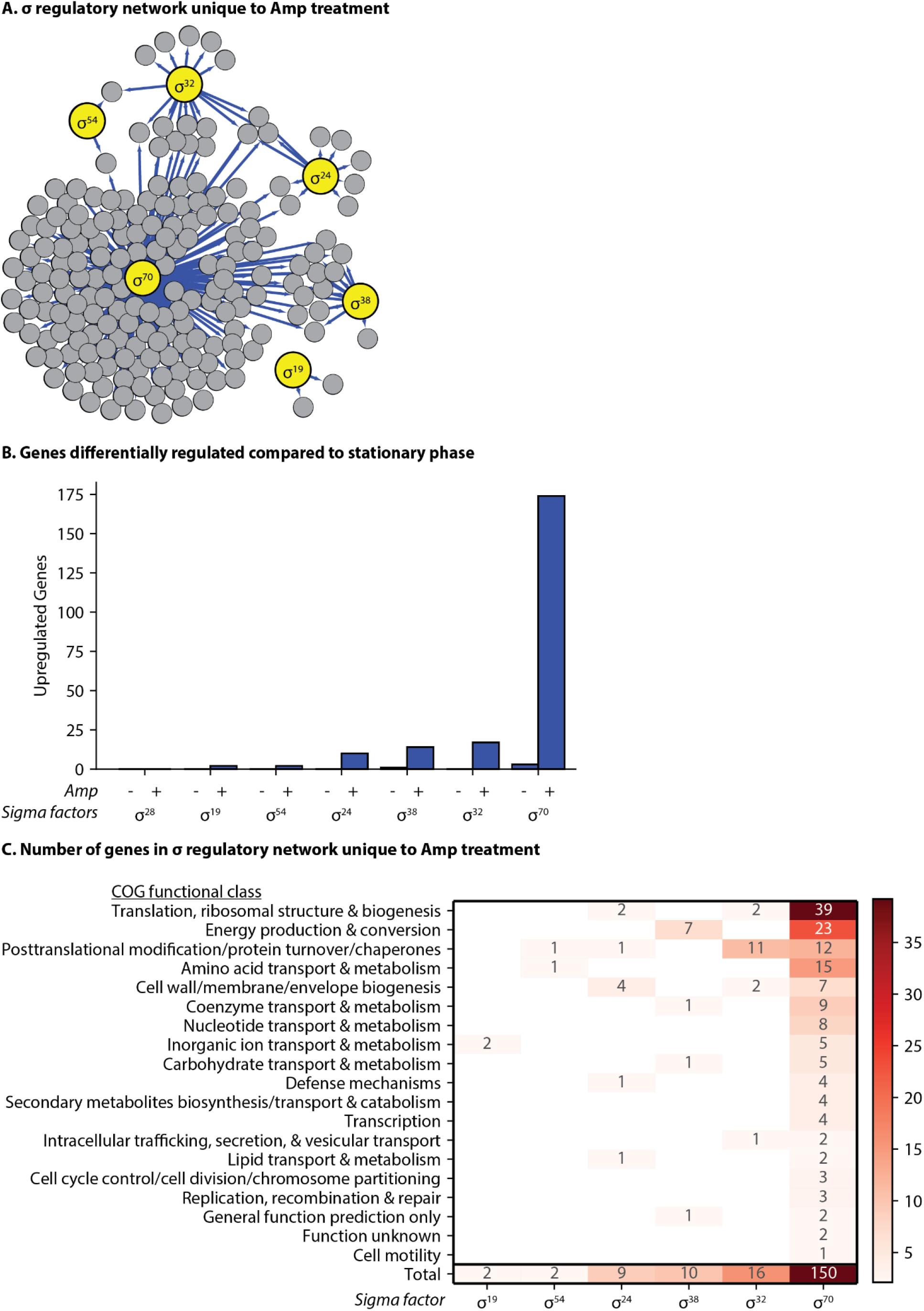
Stress related sigma factors (σ) are more active in the ampicillin treated population. Sigma factors act as global regulators that control several redundant and overlapping pathways related to antibiotic stress. Genes with increased expression over two-fold (adjusted p < 0.1) were mapped to sigma factors responsible for their transcription. **A**. A graphical representation of the sigma regulatory network during antibiotic tolerance inferred from changes in gene expression unique to the ampicillin treated population compared to stationary phase (File S2). Blue lines: increased expression. Large yellow circles: sigma factors. Small grey circles: genes regulated by the sigma factors. **B**. Counts of upregulated genes for the untreated and treated populations compared to stationary phase. Untreated (Amp-): cells incubated for 3 h in fresh media without ampicillin. Treated (Amp+): cells incubated for 3 h in fresh media with ampicillin. Genes with multiple possible sigma factors are accounted for each possible sigma factor. **C**. Changes in gene expression for each sigma factor based on COG functional group; genes uncharacterized by REF^1^ were not included.

Extensive studies on sigma factor activity, both *in vivo* and *in silico*, have identified or predicted the sigma factor(s) responsible for transcription at specific promoters throughout the genome^48–53^. We compared both the untreated and amp-treated populations to stationary phase. The total number (count) of genes that changed expression is higher in the treated population than in the untreated population as compared to the stationary phase population (Fig. 2B). As expected, the amp-treated population had more expression associated with stress related sigma factors, RpoH (σ^32^), RpoS (σ^38^), and RpoE (σ^24^), than the untreated population (p <0.001) (Fig. 2A). Many of the genes that increase in expression in response to ampicillin are regulated by both RpoD (σ^70^) as well as another sigma factor. The overlapping regulons means that some genes are accounted for twice, but the increase in expression for many of these genes is probably due to a stress-related sigma factor rather than RpoD.

As expected, there was an increase in the expression of many genes regulated by sigma factors related to stress, which have previously been associated with antibiotic tolerance^26,31^. Namely, RpoH (σ^32^; associated with heat shock^54^), RpoS (σ^38^; associated with stationary phase and stress^55^), and RpoE (σ^24^; also associated with heat shock^56^) had more active regulons in the treated compared to the untreated population. Our analysis shows that RpoH transcribes several genes related to one of the COG groups noted earlier, ‘posttranslational modification, protein turnover and chaperones’. Genes specifically regulated by RpoH encode the proteins ClpB, GroL, Lon, DnaK, DnaJ, and FtsH, many of which are related to protein degradation, which we will discuss. Meanwhile, the sigma factor RpoS regulates several genes related to energy production and conversion, and RpoE regulates a few genes related to cell wall biogenesis (Fig. 2C). While these trends are exciting to find in our data, it is clear that other components of transcriptional regulation, namely TFs, are playing a critical role in controlling gene expression and the antibiotic stress response.

### Analysis of the transcription factor (TF) regulatory network shows that the activity of global regulators increase during ampicillin treatment

TFs act on a variety of scales, from simultaneously regulating hundreds of genes (e.g. CRP^57^) to simply a few genes^58^. As with sigma factors, the activity of many TFs, especially global TFs, is regulated post-translationally (i.e. at the protein level)^57,59–61^, and therefore directly measuring expression of TFs is not typically informative in regards to their activity. We used TRN analysis to identify the role of TFs that alter gene expression in response to ampicillin.We mapped changes in expression to all possible direct regulatory interactions that could account for that change in RNA levels during ampicillin treatment. Analyzing this network makes it very clear that the regulatory pathways at play in the population’s response to ampicillin are highly interconnected and likely redundant (Fig. 3).

**Fig. 3.**
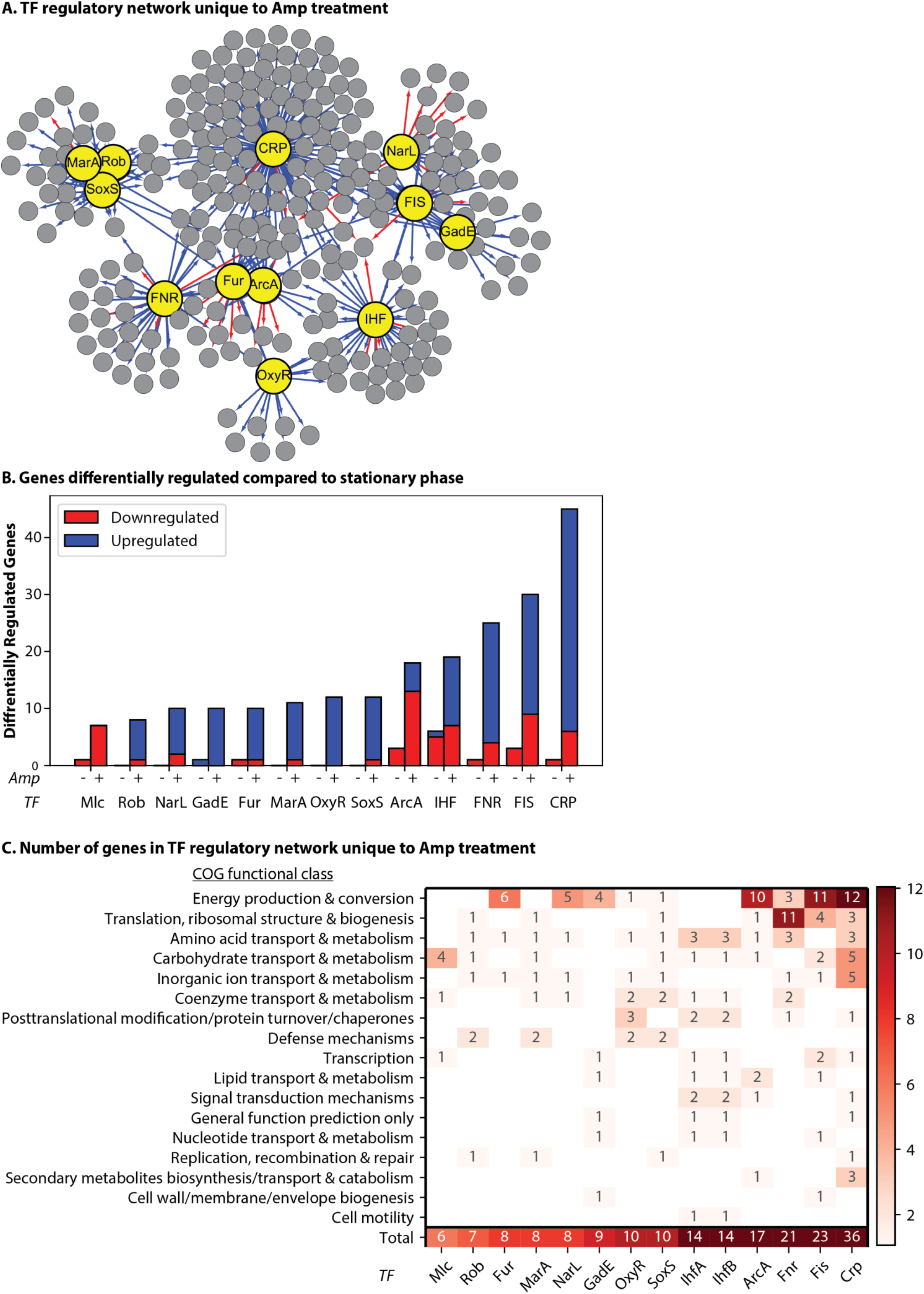
TFs associated with antibiotic tolerance are more active in the ampicillin treated population. TFs act as transcriptional regulators that control a several redundant and overlapping pathways related to antibiotic stress. Genes with changes in expression over two-fold (adjusted p < 0.1) were mapped to TFs responsible for their transcription. **A**. A graphical representation of the selected TF regulatory network during antibiotic tolerance inferred from changes in gene expression unique to the ampicillin treated population compared to stationary phase (File S3). Red lines: decreased expression. Blue lines: increased expression. Large yellow circles: select TFs. Small grey circles: genes regulated by select TFs. **B**. Counts of differentially regulated genes for the untreated and treated populations compared to stationary phase. Selected TFs displayed a significant difference (p<0.05) in the number of genes that changed expression when comparing untreated (Amp-) and treated (Amp+) groups based on the Fisher’s exact test. Amp-(untreated): cells incubated for 3 h in fresh media without ampicillin. Amp+ (treated): cells incubated for 3 h in fresh media with ampicillin. Genes with multiple possible TFs are accounted for each possible TF. **C**. Changes in gene expression for the select TFs based on COG functional group; genes uncharacterized by REF^1^ were not included. A high number of genes differently expressed are specific to a few COGs.

Several TFs are more active during ampicillin treatment, including all seven global regulators CRP, FNR, IHF, FIS, ArcA, NarL and Lrp. The impact of global TFs on the system is not entirely surprising as they are critical factors for a healthy cell physiology. Global regulators are responsible for the up and down regulation of genes across a heterogenous array of functional groups, and they are an integral facet of the hierarchical structure of the TRN in *E. coli*^*4*^. The altered activity of these TFs in the treated population suggests that the cells are actively regulating their metabolism and responding at the whole-cell level. REF^29^ found that global regulation is critical to antibiotic survival and the authors suggest that global regulators control several redundant pathways related to antibiotic stress^29^, which is also supported by our results. Furthermore, both *fnr* and *crp* have previously been identified in gene knockout studies as being related to antibiotic tolerance^62–63^.

Another global TF identified in our analysis and previously related to antibiotic tolerance is Fur. A Δ*fur* strain has been shown to have decreased levels of tolerance to ciprofloxacin but also have an increased chance of acquiring resistance^64^. Previous work examining expression data of the Δ*fur* strain with and without ciprofloxacin identified *entA, entC* and *sufA* expression as being affected by ciprofloxacin and Fur. As in, Fur regulation was critical for expression of these genes and tolerance to ciprofloxacin^64^. These three genes also show increased expression in our ampicillin treated population and have been identified in other types of tolerance as well, such as n-butanol tolerance^65^, which suggests that they are involved as a general stress response mechanism. The operon *entCDEBAH* is regulated by both CRP and Fur, encodes enterobactins, and is associated with iron homeostasis and the oxidative stress response. Moreover, enterobactins protect intracellularly against oxidative stress independently of iron availability^66^, supporting that the oxidative and antibiotic stress responses are integrated.

### Transcription factor (TF) regulatory network shows increased activity of specific, local regulators during ampicillin treatment

When we further examined pathways related to iron-sulfur (Fe-S) clusters and oxidative stress, we identified a group of genes with changes in expression regulated by local regulators OxyR and IscR. Notably, the ampicillin treated population had increased expression of genes required for Fe-S cluster formation: *sufS, sufE, sufC, iscA, iscR, iscS, iscU* and *iscX*. Fe-S clusters act as protein co-factors and serve a wide variety of functions in the cell; they are suited to a variety of processes including redox sensing, catalysis, or electron transfer. Notably, the clusters are also part of the oxidative stress response^67^. Fe-S clusters have been associated with increased development of antibiotic resistance to ciprofloxacin^64^ and uptake of aminoglycosides^68^.

Our findings further support that Fe-S clusters are involved in antibiotic tolerance. This system also highlights a restraint of our study, namely that we are only mapping regulatory events as opposed to deregulatory events. Derepression (removal of repression) is another type of transcriptional regulation that affects antibiotic survival. In this instance, when accounting for genes regulated by IscR, the *isc* genes were not included because IscR represses these genes and the genes increased in expression in our network. However, increased expression of *isc* genes is likely due to derepression of IscR, which is supported by the simultaneous increase of the *suf* genes, which IscR also regulates^67^. The same can be said for the regulation by Fur, which is also associated with iron homeostasis. While Fur was identified as significantly more active in the treated population and primarily acts as a repressor, increased expression resulting from Fur’s derepression was not accounted for in the TRN analysis. This particular group of TFs demonstrates how changes in regulatory interactions must be identified contextually with respect to when these changes might occur. The context of regulation is particularly significant for dual-regulators TFs that can activate and repress different genes depending on the state of the cell (e.g. global regulators).

Another group of local TFs that play a role in the antibiotic-treated population is the Mar/Rob/SoxS regulon. These three TFs are closely related, with degenerative consensus sequences that result in considerable overlap of regulated genes, and the regulon is positively associated with antibiotic tolerance and multidrug resistance^32–33^. Constitutive expression of SoxS has been shown to cause multiple antibiotic resistance in clinical isolates of *E. coli*^*69*^. Extensive work has shown that the TF MarA plays a role in antibiotic tolerance, as the operon was named for its role in <u>m</u>ultiple <u>a</u>ntibiotic resistance^70^. MarA is expressed stochastically and is primarily regulated by degradation at the Lon protease^71–72^. SoxS is also degraded by Lon as well as the protease FtsH. The rapid degradation of MarA and SoxS enables the cell to rapidly recover once a stress is removed^73^. We will discuss more about the importance of protein degradation to antibiotic survival later. Meanwhile, the TF Rob is in an inactive state until it is induced by binding to specific substrates^74^, a perfect example of how analyzing the mRNA level of a TF is not sufficient to understand protein activity. It is important to note that the level of *rob, marA* or *soxS* mRNA may not significantly change, while the proteins’ concentration and/or regulatory action do change. This fact underscores the importance of analyzing transcriptional networks rather than focusing on TF expression levels.

### Altered proteolytic activity affects transcriptional regulation similarly to ampicillin treatment

In addition to the regulation of MarA and SoxS through degradation, proteolytic processes play a vital role in antibiotic tolerance, as many previous works have suggested^34,75–76^. These effects on the TRN during ampicillin treatment suggest that the activity of many transcription factors is controlled at the proteolytic level and that disruption of degradation during environmental stress leads to an increase in the activity of these TFs. We have previously demonstrated that interfering with proteolytic activity causes an increase in antibiotic tolerance. To do so, we overexpressed a fluorescent protein tagged with a short amino acid degradation tag that causes it to compete for degradation at the protease complex ClpXP without affecting growth rate^34^. This competition results in a proteolytic queue, wherein proteins compete to be processed (i.e. degraded); thus the degradation of native proteins effectively slows due to an influx of new proteins with a high affinity for degradation (i.e. CFP-LAA; Fig. 4)^77–81^. The advantage of using this artificial mechanisms to slow down protease activity is it results in alteration of network dynamics without noticeable effects on growth, unlike tradition protease knockouts^34^.

**Fig. 4.**
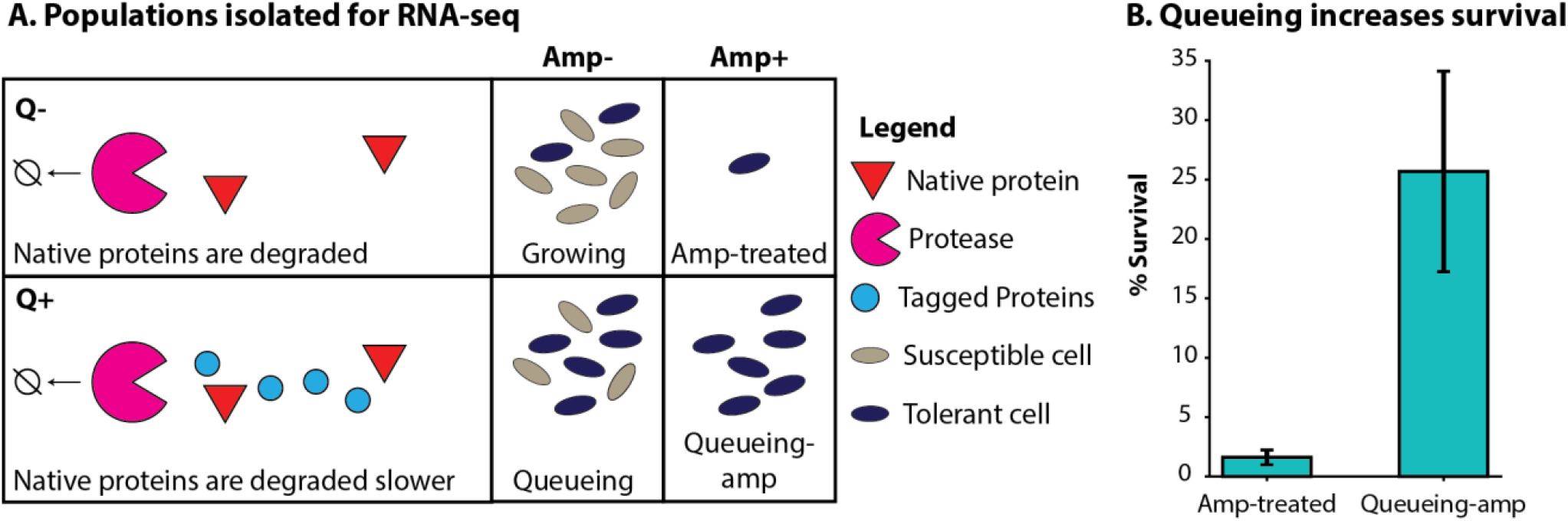
Proteolytic queueing increases survival of ampicillin. **A**. We obtained RNA-sequencing (RNA-seq) data on populations with (Q+) or without (Q-) induction of a queue at the protease ClpXP via overexpression of CFP-LAA, and populations with (Amp+) or without (Amp-) ampicillin treatment (100 μg/ml). **B)** Percent survival of cells after ampicillin treatment for three hours without a queue (Amp-treated) or with a queue (Queueing-amp).

We hypothesized that the increase in antibiotic tolerance via proteolytic queueing is due to an amplification of the transcriptional response. We compared the transcriptome of populations diluted from stationary phase into fresh media containing IPTG to induce expression of CFP-LAA. Then we incubated the cells for three hours without ampicillin (q*ueueing population*) or with lethal concentrations of ampicillin (*queueing-amp population*). We will discuss changes in gene expression as compared to stationary phase.

If slowed proteolytic activity leads to an increase in antibiotic tolerance, TRN behavior should be similar in the amp-treated and queueing populations. Indeed, that is the case. The queueing-amp population, which simultaneously responds to the induction of a proteolytic queue and ampicillin treatment, shares most of these similarities as well. The transcriptional response seen in the queuing-amp population seems to be a magnification of the response in either the queueing or amp-treated population alone, with many of the same trends (Fig. 5).

**Fig. 5.**
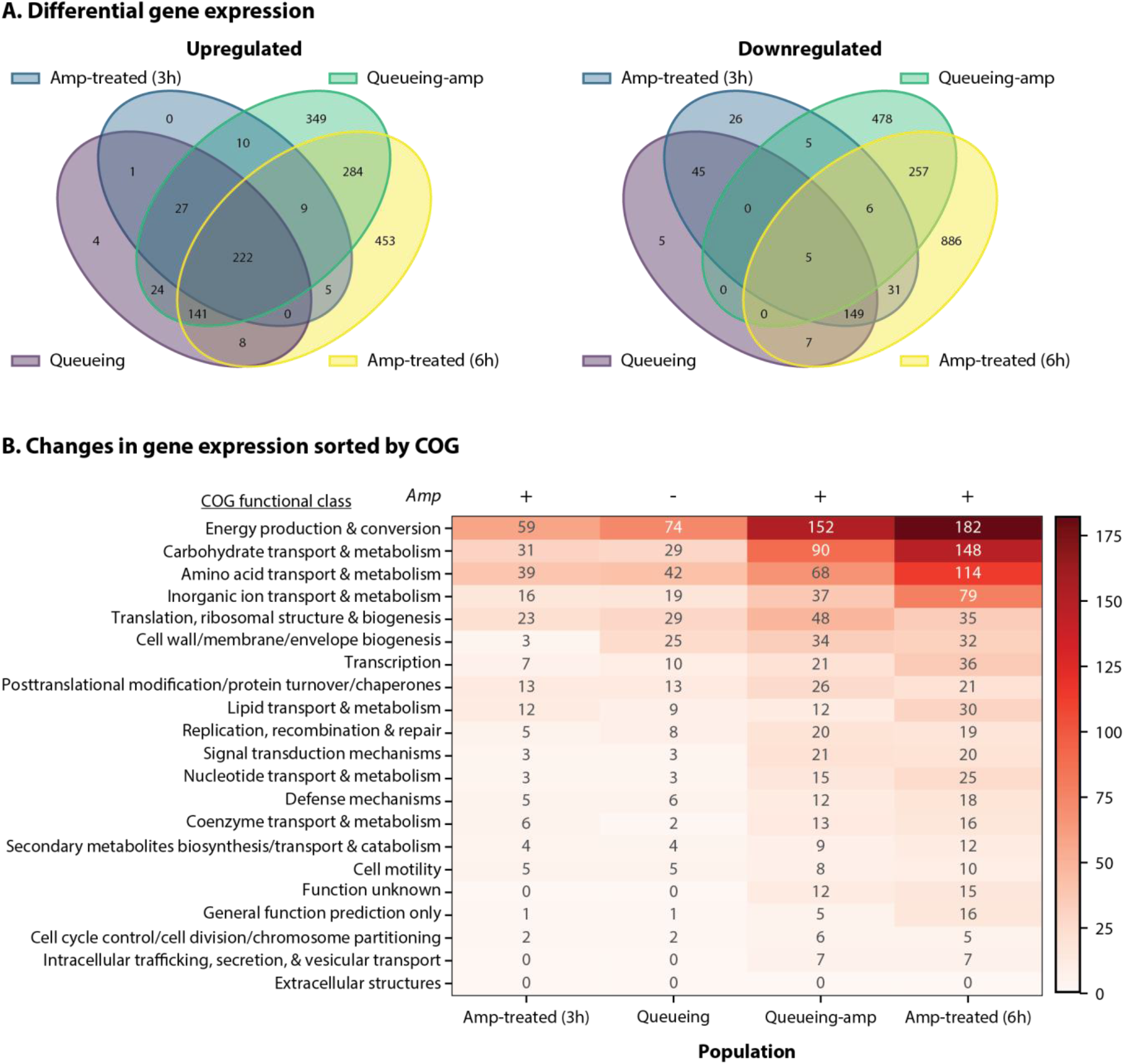
Proteolytic queueing and ampicillin treatment similarly affect gene expression in *E. coli*. Changes in gene expression compared to stationary phase (adjusted p<0.1; with >2-fold change) for each population with/without queueing (Q) and/or ampicillin (Amp). Not shown are changes in gene expression present in the untreated population. Cultures treated with ampicillin were treated for three hours (3h) or six hours (6h). **A**. Venn diagram comparing each population. Left: Upregulated. Right: Downregulated **B**. Changes in gene expression sorted in Clusters of Orthologous (COG) groups for each population. Genes in multiple categories were counted for each category.

This led us to hypothesize that the queueing is effectively ‘quickening’ (i.e. causing certain events to happen at a faster timescale) the response to the antibiotic. We then compared the ampicillin treated culture’s RNA profile after six hours to the untreated and treated populations. The gene expression profile is similar at three and six hours; however, hundreds of additional genes changed their expression by six hours (Fig. 5A). In general, the trends between gene expression in the amp-treated, queueing, and queueing-amp populations are quite similar. Upregulated genes common to the three populations include genes and systems related to tolerance discussed earlier, including those related to Fe-S clusters, oxidative stress and heat shock. When analyzing the COGs most highly regulated in queueing-tolerance, it appears that the population is heavily regulating processes related to metabolism and energy (Fig. 5B). It has previously been reported that energy levels largely affect antibiotic tolerance and persistence^26,82–83^. Particularly relevant to our work, are the findings of REF^26^, which demonstrates that lower ATP levels led to protein aggregation, which also occurs in ampicillin treated *E. coli*.

Mechanistically, protein aggregation likely results from slow protease activity, since many proteases require ATP to function^84^. In comparison, proteolytic queueing via overexpression of CFP-LAA interferes with degradation at ClpXP and causes a build-up of proteins normally degraded by the complex. The queue also affects degradation rates by other proteases^81^ and leads to protein aggregation. When looking at sigma factors and TFs that affect the transcriptional response, in this case, the alternative sigma factors are more active in the queueing-amp population compared to queueing or amp-treated cells (Fig. 6).

**Fig. 6.**
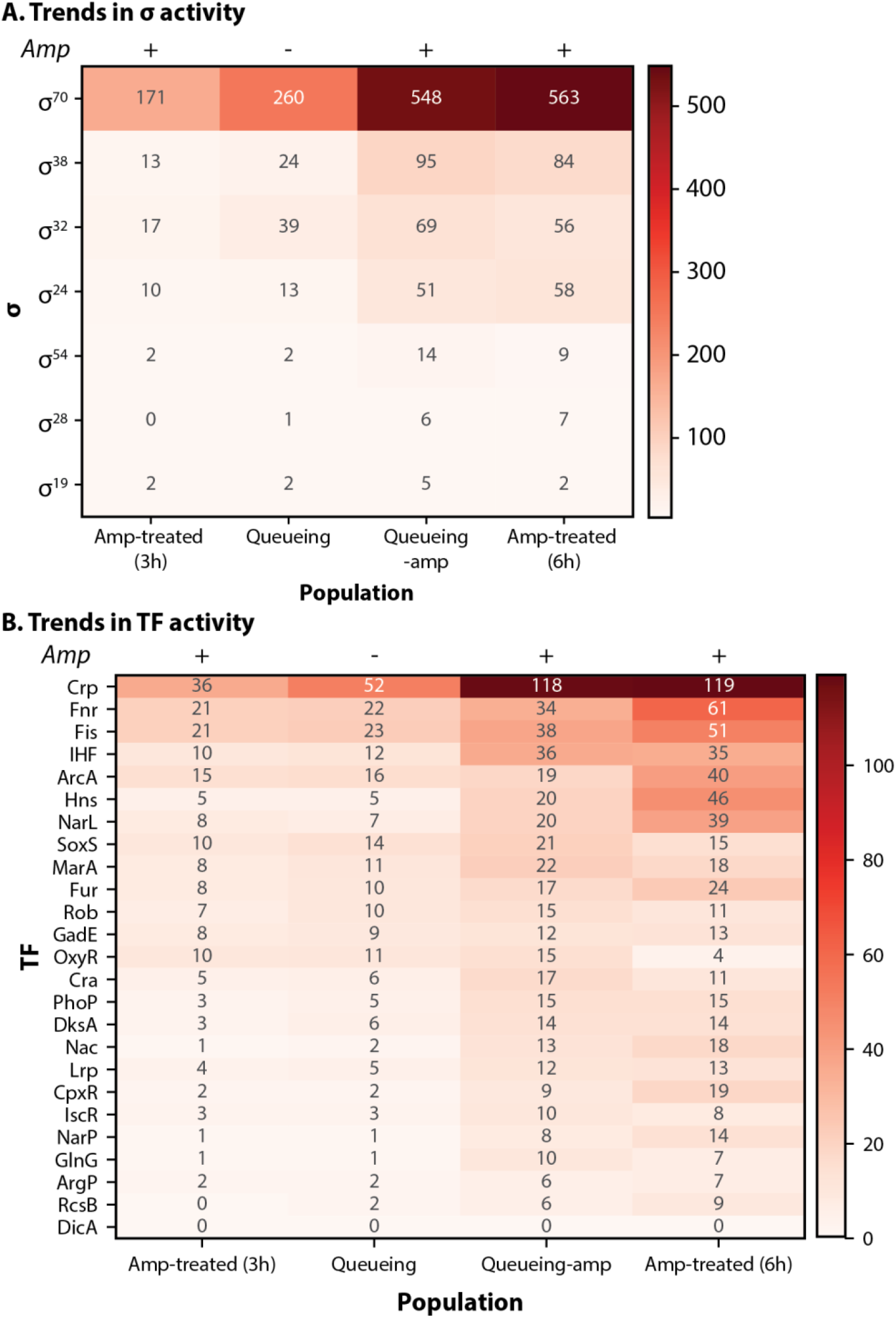
Proteolytic queueing and ampicillin treatment affect specific regulatory proteins in *E. coli*. Changes in gene expression compared to stationary phase (adjusted p<0.1; with >2-fold change) for each population with/without queueing and/or ampicillin. Not shown are changes in gene expression present in the untreated population. The amp treated population was sequenced after three (3h) and six (6h) hours of ampicillin treatment. **A**. Changes in gene expression mapped to sigma factors (σ) for each population. Genes transcribed by multiple sigma factors were counted for each sigma factor. **B**. Changes in gene expression mapped to transcription factors (TFs) for each population. Genes transcribed by multiple TFs were counted for each TF.

We propose that the magnified response in the queueing-amp population expediates the transcriptional response to the antibiotic; thus, some trends present in the amp-treated population at three hours become more apparent after six hours and/or in the queueing-amp population, which was only treated for three hours. In the case of sigma factors, we find that RpoS, RpoH and RpoE are more active in every condition compared to the untreated population (Fig. 6A).

It is important to note that some TFs that were somewhat active in the amp-treated population at three hours are now significantly engaged in the queueing-amp population and after six hours of ampicillin treatment (Fig. 6B). For example, DksA was slightly more active in the amp-treated population at three hours than in the untreated population, but it is even more active at six hours of ampicillin treatment and in the queueing-amp population. DksA is particularly interesting because it has previously been shown to play a role in antibiotic tolerance and the stringent stress response^85^, and is typically degraded by ClpXP^86–87^. The change in gene expression as the duration of antibiotic increases suggests that the cell population is altering its expression of gene networks in response to prolonged stress. It would be interesting in the future to see the expression profile with even longer exposure to antibiotics or a different class of antibiotics.

### Proteolytic degradation commonly regulates the sigma and transcription factors involved in the antibiotic stress response

As we previously touched on, many of the sigma factors and TFs involved in the antibiotic stress response are regulated by proteolysis. In the tested stressed populations (queueing and/or ampicillin), the sigma factors RpoH and RpoS are significantly more active than the untreated population based on our TRN analysis (Fig. 6A). Increased activity of these proteins is likely due to slowed proteolysis; ClpXP and FtsH rapidly degrade RpoS and RpoH, respectively^88–89^. In fact, there is direct evidence showing that ATP levels regulate RpoS degradation at the protease complex ClpXP, which slows when ATP levels are low^90^. The induction of CFP-LAA targets queue formation specifically at ClpXP, and an Δ*rpoS* strain of *E. coli* is more susceptible to antibiotics^27^. We previously attempted to replicate the queueing-tolerance phenomenon by overexpressing RpoS, but high expression levels of RpoS alone did not measurably affect antibiotic tolerance^34^. Yet, the TRN analysis of the queueing-amp population suggests that RpoS is playing a role in cell survival. This highlights the importance of looking at networks over individual genes when studying survival mechanisms to antibiotic stress.

RpoS likely works in concert with RpoH and transcription factors to mediate a response, and altered proteolytic activity could act as the coordinating factor underlying this stress response. As mentioned earlier, several genes coordinated under RpoH are related to proteolysis. Particularly noteworthy are *clpB* and *dnaK* which both have increased expression during ampicillin treatment, and have previously been demonstrated using gene knockouts in *E. coli* as critical proteins for resuscitation after removal of the antibiotic^26^. Other genes that would be key to recovering from antibiotic stress are also upregulated in the amp-treated, queueing and queueing-amp populations, such as genes for proteases *lon* and *ftsH* (both upregulated by RpoH). Increased expression of proteases likely prepares the cell to remove aggregated proteins and recover from the stress. However, both proteases are ATP-dependent; thus, their activity is regulated by the availability of ATP (especially important during recovery^26,82–83^).

### TRN analysis suggests a multifaceted response to ampicillin treatment

These results show that a coordinated transcriptional response to ampicillin results from the expression of many systems with redundant functions, as exemplified by the overlap between the Rob, MarA, and SoxS regulons. The changes we observed in the TRN under proteolytic and/or antibiotic stress indicate that cells are actively preparing for survival and recovery. The cells are likely transcribing mRNA during antibiotic treatment, especially when we consider that queueing only affects tolerance if the system is actively transcribed^34^. Furthermore, RNA levels of specific transcripts increased expression between three hours to six hours of ampicillin treatment, which supports active transcription.

We hypothesize that this transcriptional response to proteolytic queueing and antibiotic stress improve short-term survival and recovery. Especially when we consider that our previous results demonstrate that proteolytic queueing does not improve long-term survival^34^. Other factors are therefore required to survive for extended periods of time. One such possibility is translational regulation. Many previous works have demonstrated that ribosomes are largely inactive in persister cells^91–94^ resulting in slowed translation. Therefore, the transcripts we have measured may be slowly building up in the cells and are translated at low levels, if at all. The transcripts would then be readily available to the cell, and once the antibiotic is removed, ribosomes could resume protein biosynthesis. It is also possible that cells are actively translating select proteins, which seems reasonable with the number of genes upregulated during antibiotic treatment.

We also must consider that our samples are heterogeneous populations, and the data is an average of that population, which is informative but not comprehensive. Our results show that the transcriptional profile changes the more prolonged the population is exposed to antibiotics (six compared to three hours); however, it may be more complicated than that. We detected high levels of VBNCs in our populations with and without antibiotics, and our transcriptional profiles consist of a mixture of both persisters and VBNCs. Individual cells may have transcriptional profiles different from the average, and as antibiotic treatment continues, different subgroups are killed. Perhaps the surviving population after six hours of ampicillin treatment are a subpopulation present after three hours, but the expression profile is overshadowed by other subpopulations that are killed later. Understanding transcriptomic heterogeneity in the tolerant population would necessitate further experiments, such as single-cell transcriptomics, which has only recently been demonstrated in bacteria^95^. Regardless, future experiments (ribosome profiling, mass spectrometry, single-cell transcriptomics etc.) are required to understand the cellular response. Future experiments must use lethal (bactericidal) concentrations of antibiotics to draw conclusions directly related to antibiotic survival.

### Future directions to expand our understanding of antibiotic tolerance

We have demonstrated the feasibility of doing transcriptome analysis on cultures treated with lethal (bactericidal) concentrations of antibiotics and using TRN analysis to probe the network response. We used *E. coli* as our model organism because its regulatory networks are currently the most characterized of known bacteria; however, our analysis is limited to the current state of knowledge. We tested one antibiotic (ampicillin) under a single growth condition (emergence from stationary phase), and some responses we identified could be specific to these experimental conditions. Future work may be able to incorporate new information regarding the regulatory networks and contexts in which TFs and sigma factors regulate gene expression and may allow this type of analysis to be expanded to other strains of *E. coli* or different microorganisms. Exploring tolerance with a more minimal bacterial genome with less redundancy between networks (e.g. Rob and SoxS, etc.) may clarify how the TRN affects survival and recovery.

Our knowledge of the network response to antibiotics could also be expanded by using different antibiotics, with varying treatment times and growth conditions. Perhaps, the use of multiple antibiotics will lead to a greater understanding of the networks (or TFs) that are most important for survival and recovery. The recovery mechanisms after antibiotic treatment are of therapeutic interest because new drugs may be developed in combination with current antibiotics to prevent recovery, improving the effectiveness of antibiotic treatments. A recent paper highlights the potential to use combinations of antibiotics and staggered treatment of antibiotics where pairing strongly and weakly dependent metabolism antibiotics together effectively kill persisters^96^.

We have two competing hypotheses: (1) The longer the cells are exposed to lethal levels of antibiotics, the more they alter their transcriptome. (2) The longer the cells are exposed to lethal levels of antibiotics, the more they change cell types (e.g. switched from persisters to VBNCs). The ratio of VBNC to persisters may be responsible for the change in gene expression we observed. This hypothesis could be tested in the future by modifying our procedure for isolating RNA from cells treated with lethal concentrations of antibiotics, with TRN analysis of the data, and quantification of VBNC levels over time. Expanding our knowledge of antibiotic tolerance beyond *E. coli* and ampicillin treatment would significantly improve our understanding of the regulatory networks across strains and species. However, as proteolysis is a highly conserved process, it is likely that the coordination of transcriptional regulation is highly conserved during stress.

## Conclusions

Our TRN analysis and top-down approach show that studying regulatory networks in antibiotic surviving subpopulations is a powerful method to understand the molecular mechanisms that enable cells to survive lethal levels of antibiotics. Gene knockouts have been useful for identifying aspects of these networks. However, our results support that antibiotic tolerance is a whole-cell response that can occur through multiple pathways. This type of systems biology approach is advantageous compared to methods focusing on individual genes and specific mechanisms, which are susceptible to redundant systems and can often lead to contrary results. For example, toxin-antitoxins systems were once thought by many to be the primary effectors of persistence; however, several studies have suggested that they are not directly related to persistence^97–99^. We instead propose that antibiotic tolerance is a multifaceted response rooted in the general stress response systems. Indeed, in our very recent work, we demonstrated that a bacteria population containing primarily essential genes have both antibiotic tolerant and persister subpopulations to different types of antibiotics^98^. Based on ours and others’ work, we propose that a systems biology approach would be fruitful for further study of antibiotic tolerance and persistence. Many of the networks we identify and discuss in this work are universal (e.g. proteolysis, oxidative stress response), which indicates that there are likely systems common to antibiotic tolerance across several species. As such, TRN analysis of transcriptomic data is a good step towards developing a comprehensive, systems-level perspective of antibiotic tolerance and recovery from antibiotic stress.

## Author contributions

H.S.D. wrote the manuscript, performed the majority of the experiments, and ran data analysis.

T.H. performed the six hour ampicillin treatment and microscopy experiments. N.C.B. planned and directed the project. All authors contributed to discussing and editing the manuscript.

## Acknowledgements

This work is supported by the National Science Foundation award Numbers 1922542 and 1849206, and by a USDA National Institute of Food and Agriculture Hatch project grant number SD00H653-18/project accession no. 1015687. We would also like to thank Drs. Will Mather and Javier Buceta for their advice and expertise.

## Materials and Methods

### Strains and Plasmids

Strains and plasmids are from REF^34^. Briefly, the strains are derived from *E. coli* DH5αZ1 and contain the plasmid, p24KmNB82 (contains CFP-LAA under the P_lac/ara1_ promoter) as described in REF^81^. To compare results from this work to our previous work^81^, we used this same strain, and produced a queueing at ClpXP by induction of CFP-LAA. Cultures were grown in modified MMB media^34^, which consists of the following: K_2_HPO_4_ (10.5 mg/ml), KH_2_PO_4_ (4.5 mg/ml), (NH_4_)_2_SO_4_ (2.0 mg/ml), C_6_H_5_Na_3_O_7_ (0.5 mg/ml) and NaCl (1.0 mg/ml). Additionally, MMB+ consists of MMB and the following: 2 mM MgSO_4_ x 7H_2_O, 100 µM CaCl_2_, thiamine (10 µg/ml), 0.5% glycerol and amino acids (40 µg/ml). Strains containing the plasmid p24KmNB82 were grown in MMB+ kanamycin (Km, 25 µg/ml) or on Miller’s Lysogeny broth (LB) agar plates + Km (25 µg/ml). Cultures diluted after stationary phase also contained 0.1% Arabinose for consistency and CFP-LAA was induced by adding 100 μM IPTG (the promoter is P_lac/ara1_). All cultures were incubated at 37° C and broth cultures were shaken at 250 rpm.

### Quantification of antibiotic survival

Antibiotic survival was quantified as in REF^34^ with the following modifications. Cultures were grown to stationary phase in MMB+ kanamycin, then diluted 1/100 into 200 ml of pre-warmed (37°C; consistency between experiments requires the media to be prewarmed) MMB+ kanamycin+ arabinose (0.2%) and incubated in 250 ml flasks (consistency between experiments requires the use of the same flask size and media volume) for three hours. Cultures with ampicillin were treated at a concentration of 100 µg/ml. Samples were plated before and after ampicillin treatment onto LB+Km agar plates to quantify colony forming units (CFUs) and percent survival.

### Determination of cell viability

Cell viability was determined using a BacLight Live/Dead bacterial viability assay (Molecular Probes, Inc., Eugene, OR) and microscopy with a slight modification to the manufacturer’s instructions. This assay distinguishes between viable cells with intact cell membranes stained with CFDA dye (exhibit green fluorescence) and dead/dying cells with damaged cell membranes stained with propidium iodide (PI) cells (exhibit red fluorescence). In brief, 1 ml cultures (10^7^-10^8^ cells/ml) were washed with 1X PBS, 0.5 mM EDTA (pH 8.5); 3 ul CFDA (10 mg/ml) and 5 ul PI (1 mg/ml) was added simultaneously, kept in dark at room temperature for 30 min, and then subsequently washed to remove the dye. Fluorescence was measured via a Nikon Ti2E Inverted Microscope using custom scripts for image analysis. Cells were counted using “Petroff-Hausser counting chamber”. The VBNC cells was determined by subtracting the average CFU/mL from average number of live cells/ml.

### RNA isolation

Samples were treated with Tween (0.2%) and chloramphenicol (5 µg/ml) then immediately filtered out of solution (using rapid filtration similar to REFs^100–101^) and resuspended in lysis buffer containing 25 mM Tris pH 8.0, 25 mM NH_4_Cl, 10 mM MgOAc, 0.8% Triton X-100, 0.1 U/µL RNase-free DNase I, 1 U/μL Superase-In, ∼60 U/µl ReadyLyse Lysozyme, 1.55 mM Chloramphenicol, and 17 μM 5′-guanylyl imidodiphosphate (GMPPNP); lysis buffer was modified from REF^102^. Rapid filtration allows for the harvesting of any whole-cell; this minimizes RNA degradation because samples are immediately frozen in liquid nitrogen have suspension in lysis buffer. The solution was then frozen in liquid nitrogen and stored at −80°C. Frozen pellets were thawed on ice and centrifuged for 20 minutes at 16,000 *x g* at 4 °C with 0.3% Sodium Deoxychorate to pellet particulate matter. The supernatant was then removed, and then RNA was column purified (Qiagen miRNeasy cat. No. 217004) before removing rRNA (Invitrogen RiboMinus cat no. K155004) and prepping the cDNA libraries (NEBNext Ultra II cat no. E7103S). Libraries were multiplexed then pooled and sequenced by Illumina HiSeq paired-end sequencing by GENEWIZ. This isolation method is advantageous because RNA can be harvested from what are considered “viable but not culturable” (VBNC) cells and dying cells (note that all cells during antibiotic treatment are indeed dying cells, and cells only survive because we remove the antibiotic).

### DNA-sequencing

DH5αZ1 genomic DNA was harvested and purified using Genomic DNA Purification Kit (ThermoFisher) in accordance with the manufacturer’s instructions. The genome was then sequenced using paired-end Illumina sequencing Novogene Ltd. Sequencing libraries were generated from the qualified DNA samples(s) using the NEB Next Ultra DNA Library Prep Kit for Illumina (NEB, USA) following the manufacturer’s protocol. Reads were then assembled using Geneious v. 11.0.3^103^ with *E. coli* K12 str. MG1655 (NC_000913.3) as a reference sequence. *E. coli* DH5αZ1 genome is 99.9% identity to MG1655 genome, and due to the more thorough annotation of the MG1655 reference sequence, we used NC_000913.3 as the reference for RNA-seq analysis.

### RNA-sequencing data analysis

Fastq files were aligned to NC_000913.3 and the plasmid p24KmNB82 using Geneious v. 11.0.3^103^. Raw read counts were then analyzed using DEseq to compare expression (counts were normalized using the default median of ratios method); we used a 2-fold cutoff and adjusted p-values<0.1 to identify significant changes in expression^104^. All conditions tested had three biological replicates. Data has been deposited to NCBI GEO at GSE156896.

### Transcription factor (TF) network analysis

Information on transcriptional regulation was downloaded using the EcoCyc SmartTables feature^3^. We then used custom Python scripts to generate a network graph made up of directed edges toward any gene with a significant change in expression (>2-fold change in expression, adjusted p<0.01) originating from each transcription factor that may be an effector of the observed change. Visualization of this graph was done using Cytoscape^105^. The total number of genes mapped to a particular TF was compared between populations (i.e. untreated and treated each compared to stationary phase) using a two-sided Fisher’s exact test^7,106^.

### COGs

Data was downloaded from http://www.ncbi.nlm.nih.gov/COG/^1,45^.

## Data and materials availability

All data that supports the findings of this study are available from the corresponding author upon request.

## Supplemental information

File S1: Tab-delimited file containing direct regulatory information for each gene according to the Ecocyc^3^ database as of February 2020.

File S2: Tab-delimited file containing regulatory interactions used to make network in Fig. 2A

File S3: Tab-delimited file containing regulatory interactions used to make network in Fig. 3A

